# A model of the PI cycle reveals the regulating roles of lipid-binding proteins and pitfalls of using mosaic biological data

**DOI:** 10.1101/2020.05.26.116251

**Authors:** Francoise Mazet, Marcus J. Tindall, Jonathan M. Gibbins, Michael J. Fry

## Abstract

The phosphatidylinositol (PI) cycle is central to eukaryotic cell signaling. Its complexity, due to the number of reactions and lipid and inositol phosphate intermediates involved makes it difficult to analyze experimentally. Computational modelling approaches are seen as a way forward to elucidate complex biological regulatory mechanisms when this cannot be achieved solely through experimental approaches. Whilst mathematical modelling is well established in informing biological systems, many models are often informed by data sourced from different cell types (mosaic data), to inform model parameters. For instance, kinetic rate constants are often determined from purified enzyme data *in vitro* or use experimental concentrations obtained from multiple unrelated cell types. Thus they do not represent any specific cell type nor fully capture cell specific behaviours. In this work, we develop a model of the PI cycle informed by *in-vivo* omics data taken from a single cell type, namely platelets. Our model recapitulates the known experimental dynamics before and after stimulation with different agonists and demonstrates the importance of lipid- and protein-binding proteins in regulating second messenger outputs. Furthermore, we were able to make a number of predictions regarding the regulation of PI cycle enzymes and the importance of the number of receptors required for successful GPCR signaling. We then consider how pathway behavior differs, when fully informed by data for HeLa cells and show that model predictions remain relatively consistent. However, when informed by mosaic experimental data model predictions greatly vary. Our work illustrates the risks of using mosaic datasets from unrelated cell types which leads to over 75% of outputs not fitting with expected behaviors.

**Authors summary:** Computational models of cellular complexity offer much in terms of understanding cell behaviors and in informing experimental design, but their usefulness is limited in them being built with mosaic data not representing specific cell types and tested against limited experimental outputs. In this work we demonstrate an approach using quantitative proteomic datasets and temporal experimental data from a single cell type (platelets) to inform kinetic rate constants and protein concentrations for a mathematical model of a key signaling pathway - the phosphatidylinositol (PI) cycle; known for its central role in numerous cell functions and diseases. After using our model to make novel predictions regarding how aspects of the pathway are regulated, we demonstrate its versatile nature by utilising proteomic data from other cell types to generate similar predictions for those cells while highlighting the pitfalls of using mosaic data when constructing computational models.

## Introduction

The phosphatidylinositol (PI) cycle is a key component of the signaling machinery downstream of receptor protein-tyrosine kinases (RTK) and G protein-coupled receptors (GPCR). The cycle can be found in all eukaryotic cells, is the source of multiple second messengers through the actions of phospholipase C (PLC) and phosphoinositide 3-kinase (PI3K) and is assumed to function the same way in different cell types. The universal nature of the pathway means it is of wide interest, but its multiple components are technically difficult to measure, making it a good candidate to explore using mathematical modelling approaches. There has been a number of prior attempts to model aspects of the PI cycle looking at portions of the signaling cascade using ordinary differential equations (ODEs), informed to a large degree by data from different cell types [1–3]. We have been unable, however, to combine them into a single model of the complete pathway and recapitulate the different published biological outputs. This issue applies to a number of cell signaling models, with their development being hampered by a lack of cell type specific biological data to inform kinetic rate constants and protein concentrations. In particular, we note current cell signaling pathway models often lack in-vivo cell-specific time course data to inform model parameter values. They also usually incorporate purified enzyme kinetic data which may bear little resemblance to the *in-vivo* kinetics [4], or experimental values obtained from multiple, unrelated, cell types, with reactant concentrations often estimated. Signaling can be cell context dependent producing specific responses to stimuli and “mosaic” models. Although useful in investigating and informing the general biological processes involved, models informed in these ways may not necessarily recapitulate cell specific dynamic behaviors.

We postulate that cell-type specific datasets generated by omics approaches coupled to time-course analysis of the gene expression or protein modifications would provide a more consistent approach to informing mathematical models. This would allow us to focus on determining *in-situ* reaction kinetic rate constants which once obtained should be possible to use for other cell types as long as specific quantitative proteomic data are available. Here we describe a PI cycle model making use of a quantitative proteomic dataset [5] and experimental phospholipid (PL) and inositol phosphate (IP) time-course data produced in platelets, under similar conditions (Fig 1A) [6–11]. We use the model to demonstrate the importance of lipid-binding proteins in regulating homeostasis and to inform the regulation of several key proteins in platelets. We next investigate how the model can be used to simulate the PI cycle in other cell types. In doing so we leave our rates unchanged but inform the model with specific cell-type proteomic data (Fig 1B). Finally, we demonstrate the pitfalls of using mosaic data to inform such a model and how it can lead to erroneous conclusions. We discuss how this limits the use of combining cell signaling models and directing the design of future experiments.

**Fig 1.**
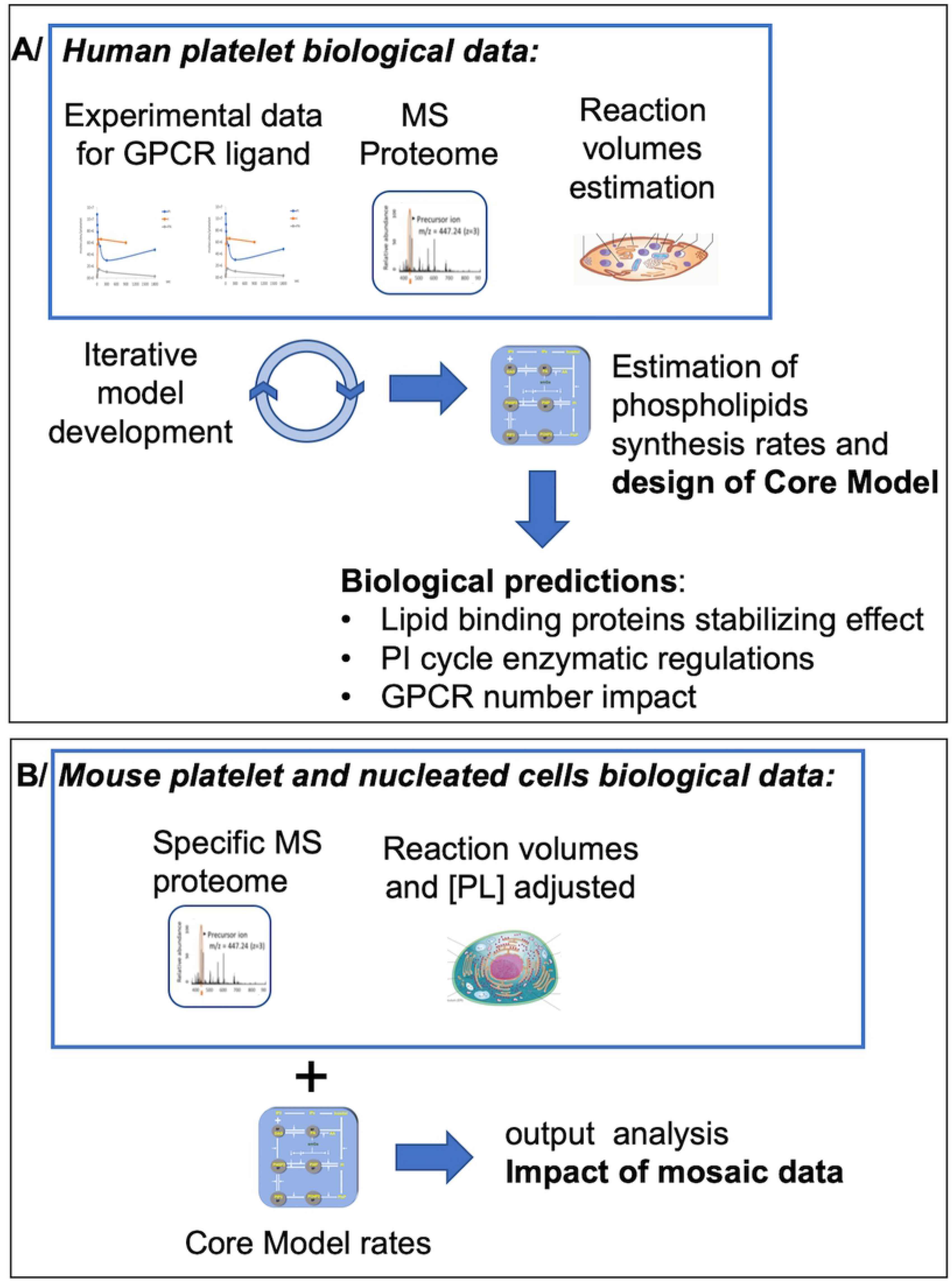
Graphical summary of the different modelling steps. A: The core model was produced using proteomics and signaling data (obtained with a single GPCR ligand, Thrombin) obtained for human platelets and used to generate predictions regarding PI cycle driven cell signaling. B: mouse platelet and nucleated cell proteomic data (Cell type B) were used to populate our Core Model, generate output predictions and analyse the impact of using data from different origins in a single model.

## Results

We first sought to combine previously published mathematical models of the PI pathway in order to construct a model of the pathway in platelets. Whilst the structure of the pathways and their respective mathematical formulations were generally similar, how the respective reaction rate constants and concentrations were informed varied greatly. For instance, values were not consistently reported both in terms of their magnitude and units, it was not always clear how all values had been informed and values to inform cell type specific models had been obtained from data related to different cell types. We also sought to consider a previous model of PI signaling in platelets [1], which we could extend to account for our needs. We found that whilst the model had been useful in informing platelet biology, it was informed using a range of different cell type data and could not meet our needs.

In light of these points and knowing that considerable biological data for informing platelet biology is now available, we thus decided to formulate a complete model of the PI cycle in platelets, exclusively using specific parameters for PI cycle pathway proteins, phospholipid substrates and resulting second messengers (Fig 2). In respect of platelet specific data, literature mining revealed three quantitative proteomes for human and mouse platelets and HeLa cells that contained data on all the key proteins required for the PI cycle to function [5,12,13]. Further literature mining revealed that detailed sets of time-resolved data on complexes formed in the PI cycle pathway were also available for human platelets (Fig 3A, 3C, S1 Fig). Using this information, we first developed a human platelet “Core Model” that focuses on G protein-coupled receptor (GPCR) signaling through Gαq, leading to PLCβ activation and production of inositol 1,4,5-trisphosphate (IP3) second messenger. Phosphatidylinositol (3,4,5)-trisphosphate (PIP3), produced via Gαi and PI3K, and other PIP3-derived PL were also monitored, but their highest levels were systematically two orders of magnitude lower than PL and Inositol (Ins) [9,14], (Fig 2A, S1 Fig, S1Table). As such we assumed they are unlikely to significantly alter the PLCβ results and conclusions and subsequently were excluded in the model we developed.

**Fig 2.**
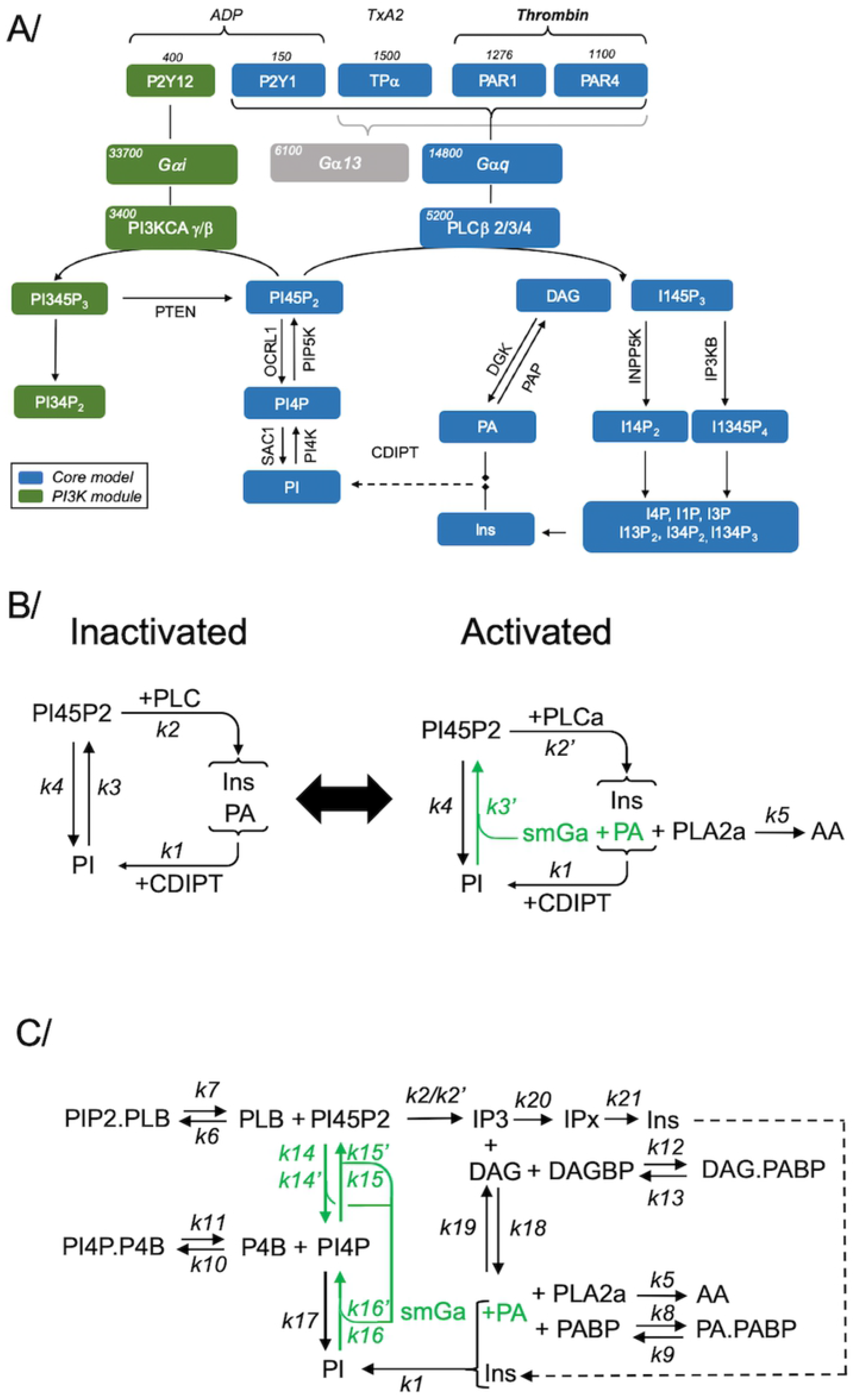
Schematic illustration of the PI cycle in mammalian cells and model iterations. A: Schematic of the GPCR ligands and reactions in platelets covered by our model (blue). PI3K-dependent (green) and Gα13-dependent (grey) reactions were not included in the core model. Abbreviations: Phosphatidylinositol (PI), Inositol (Ins), Phosphatidylinositol-3,4,5-trisphosphate (PIP3), Phosphatidylinositol-4,5-bisphosphate (PI45P2), Phosphatidylinositol-3,4-bisphosphate (PI34P2), Phosphatidylinositol-4-phosphate (PI4P), Inositol trisphosphate (IP3), Diacylglycerol (DAG), Phosphatidic Acid (PA). B: Early iterations used a simplified Core Model to estimate the phospholipids synthesis rates in inactivated cells (+PLC) and activated cells (+PLCa). Rate labels are indicated as either basal rates (*k*) or activated rates (*k’*). C: The completed Core Model schematic including lipid binding proteins.

**Fig 3.**
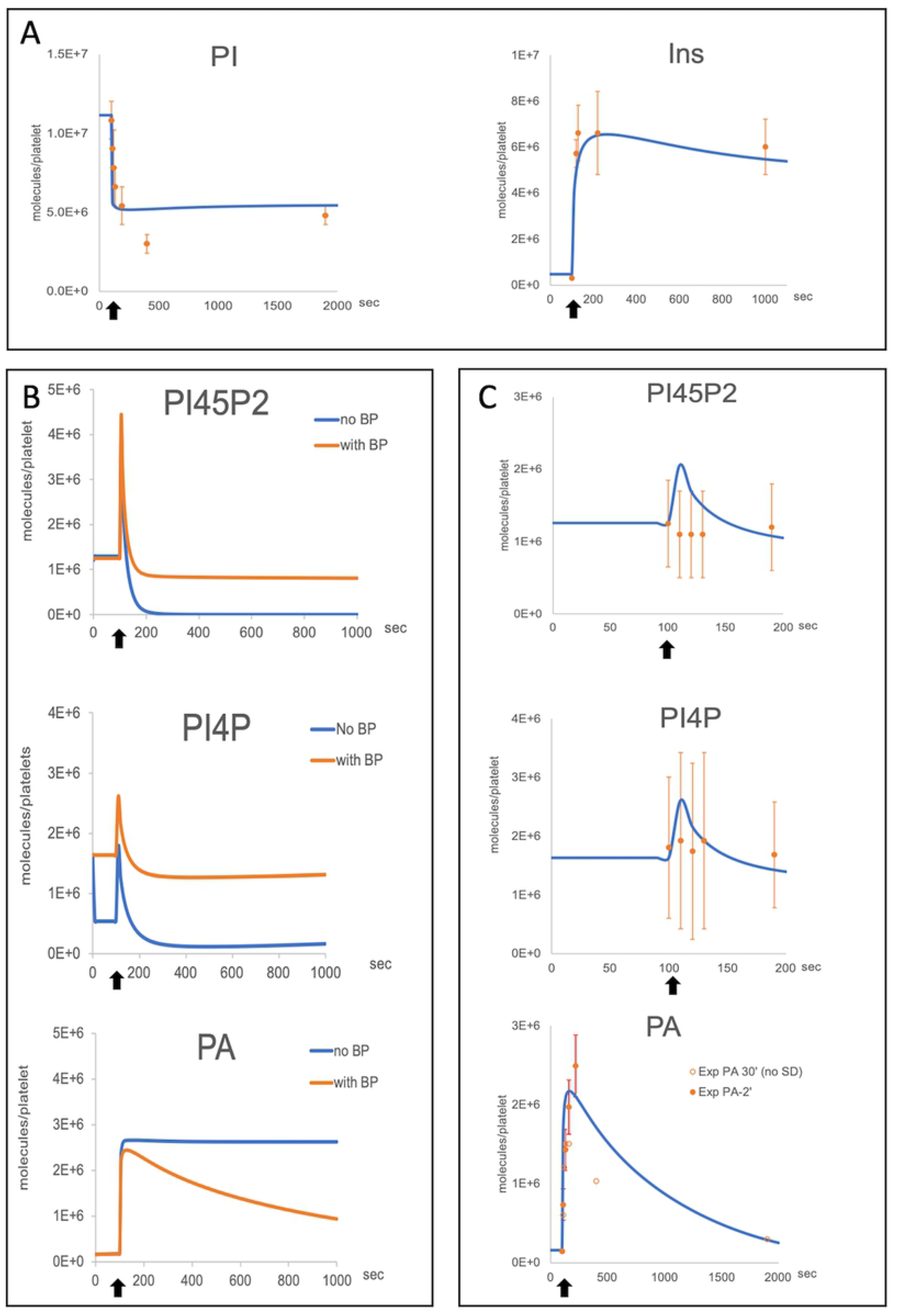
Comparison of experimental results and model simulations. A: PI and Ins experimental results (orange dots with SD) and simulations (blue curves) after Thrombin stimulation. Secondary signaling by other, secreted, GPCR ligands is taken into account and assumed to happen immediately. B: Early model iterations simulations showing the stabilizing impact of lipid binding proteins (BP) on the dynamics of PI45P2, PI4P and PA before and after activation with Thrombin. The numbers of BP and the ON/OFF rates of their binding to their relevant PL were determined using parameter scans. C: The final simulations in our completed core model are shown together with the experimental results. Experimental results are shown with standard deviations when available. (References, SD and model parametisation procedures in Methods and Supplementary Informations). Experimental data: n=10, SEM are shown. Activation is indicated by arrow.

### The Core Model – The human platelet model

We constructed an ODE model of the PI cycle in human platelets based on the model shown in Figure 2A, which was solved in COPASI (v4) [15]. Full details on how the model was developed, parameterized and solved are presented in the Supplementary Information. Briefly, in order to accurately inform the full model parameterization and given the full model consisted of 35 parameter values, we decided to use an iterative approach of reduced models to both decrease the size of the parameter space being determined, whilst increasing confidence in the parameter values determined. We started with the most simplified model shown in figure 2B, and via model-data fitting using the parameter estimation algorithm available in COPASI, used this to inform the relevant parameter values. This model was then extended in the next iteration with a number of additional reactions, which were chosen so as to not greatly increase the size of the undetermined parameter set. Each model was informed where possible with the previously determined values, whilst new unknown parameter values were again determined. Iterative steps in building up the complexity of our model in this way, allowed us to inform the full model pathway as detailed in Table S5, S1 Fig and the Supplementary Information.

### Phospholipid binding proteins stabilize phospholipids variations

An important point that arose during the development of the Core Model, was that solely adapting the rates of production or recycling of different PL to switch from non-actived to activated states, and back again did not lead to a biologically realistic solution. We concluded that the problem was linked to the availability of PL to the enzymes in addition to the rates at which they were being used in the cycle. It is known that PI4P and PI45P2 do not alter dramatically upon stimulation from their homeostatic levels [2,6,16,17]. We hypothesized that this might be due to the presence of PL binding proteins that would sequester membrane phospholipids through protein-lipid interactions, which have been shown to be important for numerous cell functions [18].

We first focused on PI45P2-binding proteins and added these to our model with reversible binding reactions with PL to control its use by PLCβ (Fig 2C, S1 and S2 Figs.). Parameter scans of the amount of binding proteins and their binding rates predicted that to correctly simulate temporal PI45P2 changes, the number of binding proteins should be around 1.3 x10^6^ per cell. A detailed search for known PI45P2-binding proteins in the proteome dataset [5] revealed, in close agreement with our prediction, a value of 1.12 x10^6^ per cell (S2 Table). Similar methodologies were used for PA, PI4P and DAG, and led to model results matching the experimental data (Fig 3B-C, S2 Fig).

### Model predictions of the regulation of PI cycle enzymes

PI45P2 and PI4P homeostatic levels in human platelets are similar, at around 1.2-1.8 x 10^6^ molecules per cell respectively, while the estimated amount of plasma membrane PI is around 6 x 10^6^ molecules per cell (S1 Table). This suggests unbalanced phosphorylation/dephosphorylation reactions between PI and PI4P and balanced reactions between PI4P and PI45P2 in inactivate cells. The addition of PI4P, its binding proteins and its metabolizing enzymes PI4K, OCRL1 and SAC1, could only achieve the correct dynamics of PI4P, PI45P2 and PI observed both before and after activation when the rates of the PI4K and OCRL1 were regulated in a manner similar to PIP5K i.e. all three enzymes have similar low kinetic levels before activation, which are increased upon GPCR activation. In contrast, the SAC1 kinetic rate constant needed to remain unchanged after GPCR activation to recapitulate the correct PL dynamics.

In conclusion, our results suggest that the regulation of the PI cycle in both inactive and activate cells is achieved by mechanisms differentially controlling the enzymes involved. PI4K and PIP5K have been suggested to be scaffolded by proteins at the membrane, exchanging PL almost directly [19]. Our simulations suggest that OCRL1 is also part of this complex or co-localises in the plasma membrane, being then regulated in concert with the kinases. SAC1, however, is regulated differently than the other PI cycle enzymes which leads to the hypothesis that it may not co-localize with them.

The final step of our model development was to reintroduce PLCβ products, IP3 and DAG, as intermediates for Ins and PA. While DAG and PA levels, regulated by two simple reactions and their respective binding proteins could be easily modeled, cytosolic IP3 and Ins levels could not be correctly simulated by a single direct reaction. We were only able to match their respective dynamics by adding an intermediate step which removes IP3 rapidly, reflecting the production of IP4 by the kinase IP3Kb and IP2 by the PL phosphatase INPP5. This is followed by slower production of Ins by a complex set of reversible reactions which we simplified in our model and wrote as a single reaction (Fig 2A and 2C, S1F Fig) [20].

### Gαq-coupled receptor number governs the strength of IP3 production

We chose to model the PI cycle because of its central role in signalling cascades triggered by a variety of GPCR agonists. One issue with this approach is that the strength of the signalling is highly variable depending on the type of receptor involved, with some researchers arguing that differences in molecular identity, biological activation processes and post-activation recycling of the receptors are responsible for this variability. Kinetic studies, however, demonstrate that GPCR signalling is primarily and rapidly down-regulated at the level of the receptors by phosphorylation, and by inactivation of their direct partners, the G proteins, and the RGS and PLCβ proteins [21,22]. Together the evidence suggests that the strength of the GPCR signalling is actually a function of receptor abundance [1,23].

The experimental data we used to inform our model kinetic rate constants were obtained after Thrombin activation of the platelets. However, the experimental methodologies used by the different authors suggest that secondary GPCR signalling through secreted ADP and TxA2 release was also activated [6–11,16]. We originally designed our Core Model to account for the activation of all their respective Gαq-coupled receptors, namely PAR1/4 for Thrombin, P2Y1 for ADP and TPα for Thromboxane (Fig 4A-B, RGq =5000, S4 Table). By solely reducing the receptor numbers to simulate only ADP activation via the P2Y1 receptor (Fig 4B, RGq =150), we were able to simulate the experimentally observed IP3 output supporting the claim that, in the case of Gα-coupled receptors, the strength of the signalling is indeed related to the number of receptors being activated [24] (Fig 4B, S4 Table).

**Fig 4.**
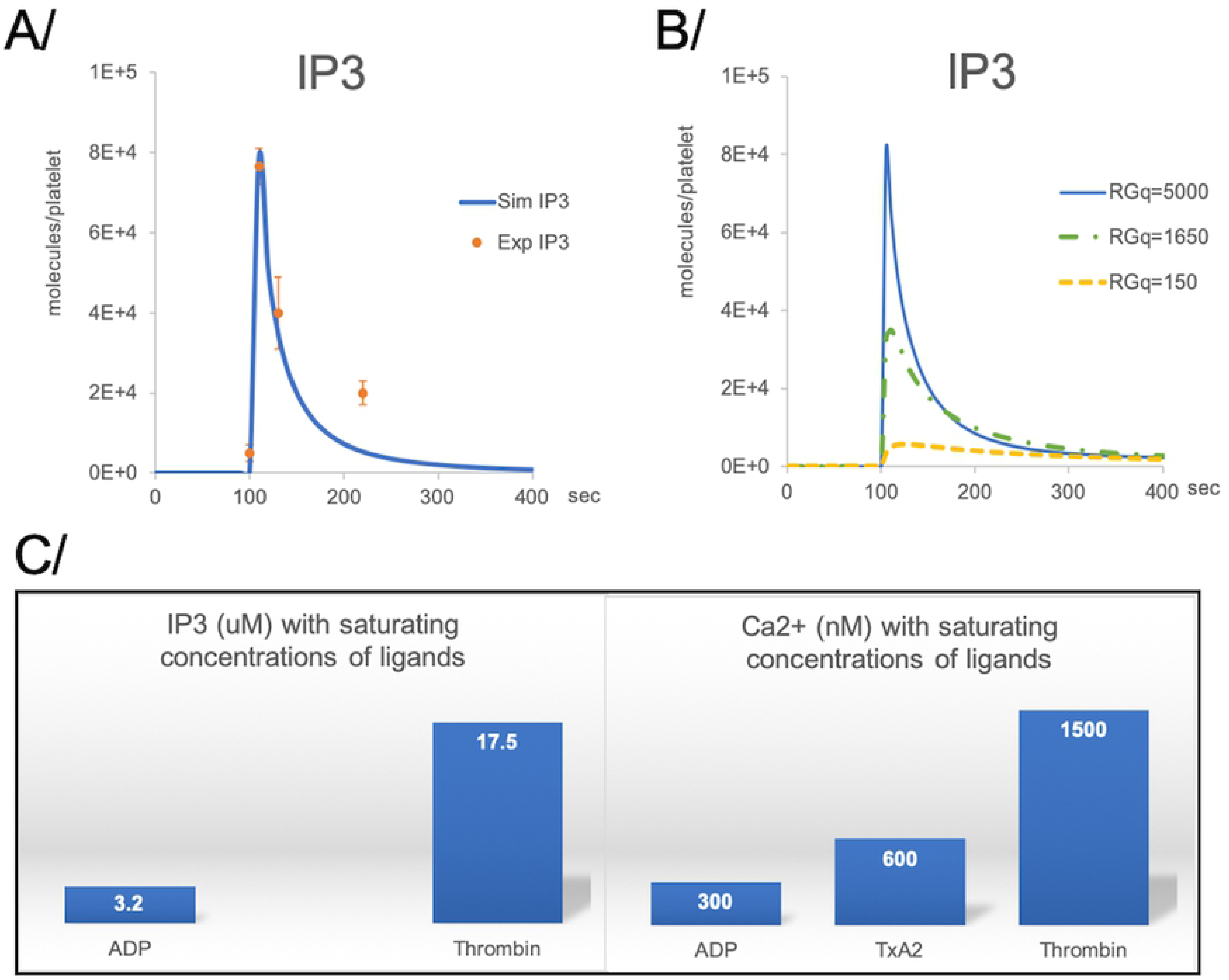
IP3 simulations with differential receptor numbers. A: IP3 experimental results (orange dots with SD) and simulations (blue curves) after Thrombin stimulation. B: simulations of the impact of Gαq-coupled receptor numbers (RGq) on IP3 production. RGq numbers reflect primary and secondary GPCR ligand activation (Thrombin followed by TxA2 and ADP: RGq = 5000; TxA2 followed by secreted ADP: RGq= 1650; ADP alone: RG1=150). C: comparison of known IP3 and calcium mobilization results in platelets. in absence of experimental data on IP3 production after TxA2 activation, we are using known cytosolic calcium concentrations as a proxy [25]. The simulations of IP3 release from a receptor number corresponding to a TxA2/ ADP activation (RGq = 1650) show an intermediate response as is also seen for calcium mobilization experimental data.

We could not find data regarding IP3 levels after TxA2 activation, however, IP3 triggers calcium cellular mobilization and both experimental and mathematical models have shown a relationship between cytosolic IP3 and calcium levels [17,26]. Quantification of calcium in platelets shows that TxA2-mediated activation leads to half of calcium being mobilized compared to Thrombin. ADP activation only triggers a fraction of IP3 and calcium release compared to Thrombin and TxA2 [25,27] (Fig 4C). Interestingly, the number of TPα and P2Y1 receptors which would be involved in a TxA2 primary activation followed by the secreted ADP secondary activation is just under half the number of the full Gαq-coupled receptor complement (S4 Table). When modeling the likely IP3 output following the activation of TPα and P2Y1 receptors, we obtained a predicted peak value for IP3 roughly half that obtained with Thrombin (Fig 4B, RGq=1650). This supports the hypothesis that IP3 levels regulate the intensity of calcium release in a GPCR receptor number dependent manner.

### Applying the Core Model to other cell types

Given the universality of GPCR signaling and the PI cycle in mammalian cells we assume that the network structure and kinetic rate constants do not vary greatly between platelets and other cell types. In addition, IP3 dynamics and PIP2 stability have been described in other cell types and show comparable behaviors to those observed in platelets [2,17]. We thus hypothesised that we should be able to simulate the PI cycle and IP3 production in other cells and obtain similar output dynamic patterns by using cell specific protein initial concentration values, whilst leaving our model structure and kinetic rates unchanged (Fig 1B).

#### Application to the mouse platelet using proteome data

Before proceeding to consider how applicable our model was in nucleated cells, we first used data from the mouse platelet proteome, to check the model behavior [12]. We corrected for the difference in size of the reaction compartments (25% of those in human platelets, S4 Table) but left the ODEs untouched. Despite the protein concentrations being sometimes very different from their human counterparts, the simulations show almost identical temporal behaviors of the different PL and IP as for human platelets and levels in line with the initial PL abundancy (S3A Fig).

#### Application to nucleated cells using cell-specific data

To demonstrate the wider utility of our model we next created a generic simulated nucleated cell, using an average volume of 2000 fl and calculated the size of the reaction compartments as described earlier. PL, IP and lipid-binding protein numbers were adapted from the human platelet data (S4 Table). We first simulated the behavior of an enlarged platelet by populated our nucleated PI cycle model with the protein copy numbers of a human platelet scaled to match the new reaction volumes (labelled pltx17). Next, protein copy numbers from the epithelial adenocarcinoma HeLa human cell line proteome dataset were used to populate the same model [13]. In order to compare the results and in the absence of common data regarding GPCRs, the activation in these simulations were performed using 85000 molecules of GPCRs, i.e. corresponding to the same concentration of receptors for a normal human platelet. Given that HeLa cell protein numbers are drastically different from the enlarged platelet simulation, we expected a radically different series of outputs. HeLa cell simulations did indeed show some differences in the peak concentrations of the PLs and IPs we surveyed compared to the large platelet simulation, but overall the temporal dynamics were similar (Fig 5), suggesting our model solutions are robust to changes in protein concentration and exhibit the correct PLCβ-dependent IP3 signaling response.

**Fig 5.**
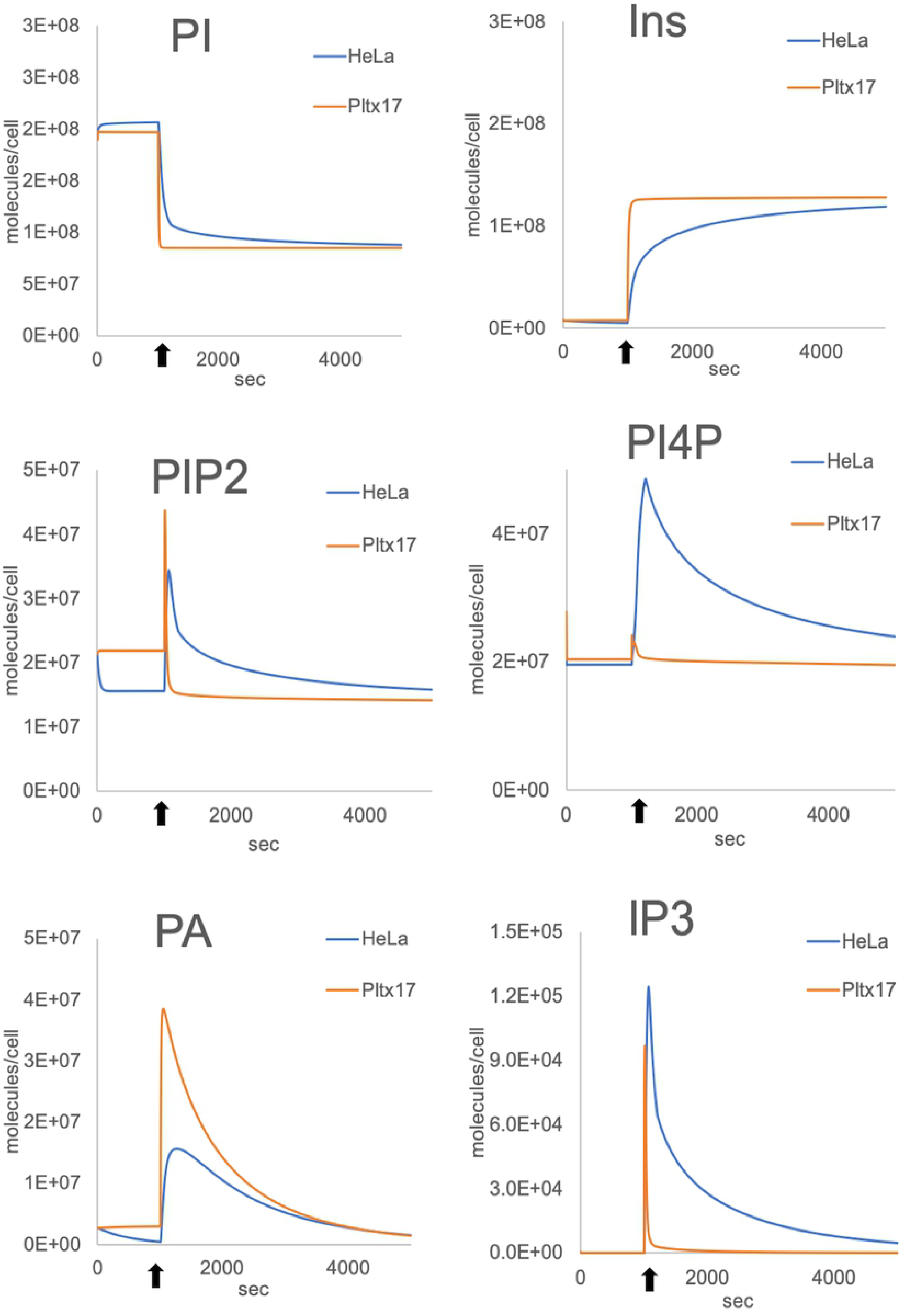
Application to other cell types. A: GPCR-PLCβ simulations using Hela proteome dataset (dashed line, (12)) and compared to a hypothetical large platelet (pltx17, continuous line). Reactions and kinetic parameters are unchanged from the original core model; reaction volumes, initial PL numbers and their binding proteins were adapted as described in Methods and Table S4. Simulations are run for 5000 seconds with activation occurring at 1000 sec (arrows).

### The effect of using mosaic proteomic datasets for informing model parameters

After demonstrating the portability of our PI cycle pathway model to other cell types and its robustness when parameterised with cell specific protein numbers, we wanted to investigate its response when informed by a mosaic dataset. Using our nucleated cell model, we used a proteomic dataset from the bone osteosarcoma epithelial U2OS human cell lines with only partial data regarding the PI cycle enzymes and missing the concentration of Gαq, IP3 processing enzymes (IP3Kb and INPP5), OCRL1, PI4K and cPLA2 proteins [28] (S4 Table). HeLa protein concentrations were used to inform the missing values. The resulting simulations led to PI, Ins and PA concentration profiles similar to the Hela simulations although the concentration of PI45P2, PI4P and IP3 were much lower than expected for a cell of this size. IP3 production is significantly affected and unlikely to lead to a realistic outcome (Fig 6). We then utilised a series of parameters scans, to inform those protein concentrations for which values were not available. Whilst we were able to find protein copy numbers which meant Core Model simulations could replicate the previously modelled HeLa cell behaviour (S4 Table, Fig 6), the use of partial data from a different cell type would not necessarily provide the correct outputs.

**Fig 6.**
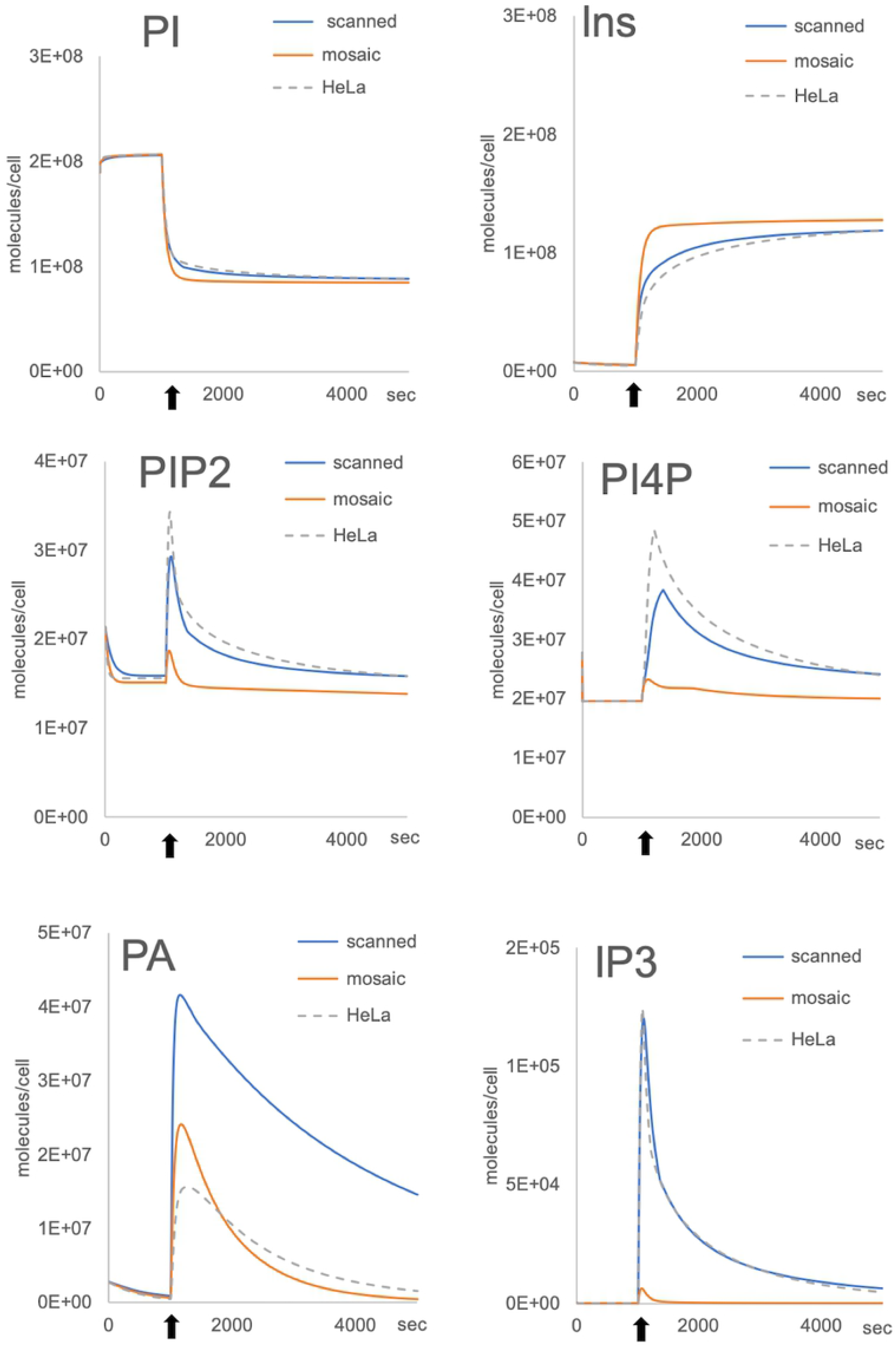
Simulations with mosaic proteome datasets. The values for PI4K, OCRL1, Gαq, cPLA2 and IP3 modifying enzymes are missing from the U2OS proteome. We performed a series of scans to estimate the missing protein numbers (“scanned”) that allow results similar to those obtained with the HeLa dataset (“HeLa”) and compared them to a mosaic dataset created by using HeLa protein numbers to replace the missing U2OS numbers (“mosaic”). The results show that the mosaic dataset generates outputs for PI45P2 (PIP2), PI4P and IP3 are unlikely to lead to a cell signaling response.

We then considered random combinations of all the protein concentrations from the U2OS and HeLa datasets. The simulations showed no consistency for any PL or IP we surveyed, with 75% leading to incorrect behaviours (Fig 7). Whilst 25% of the results produced the correct behaviour for IP3, these simulations did not always lead to correct outputs for the other PLs we surveyed such as PI4P or PA. Ultimately this suggests that combinations of protein concentration values may lead to incorrect model approximations of the underlying protein concentrations.

**Fig 7.**
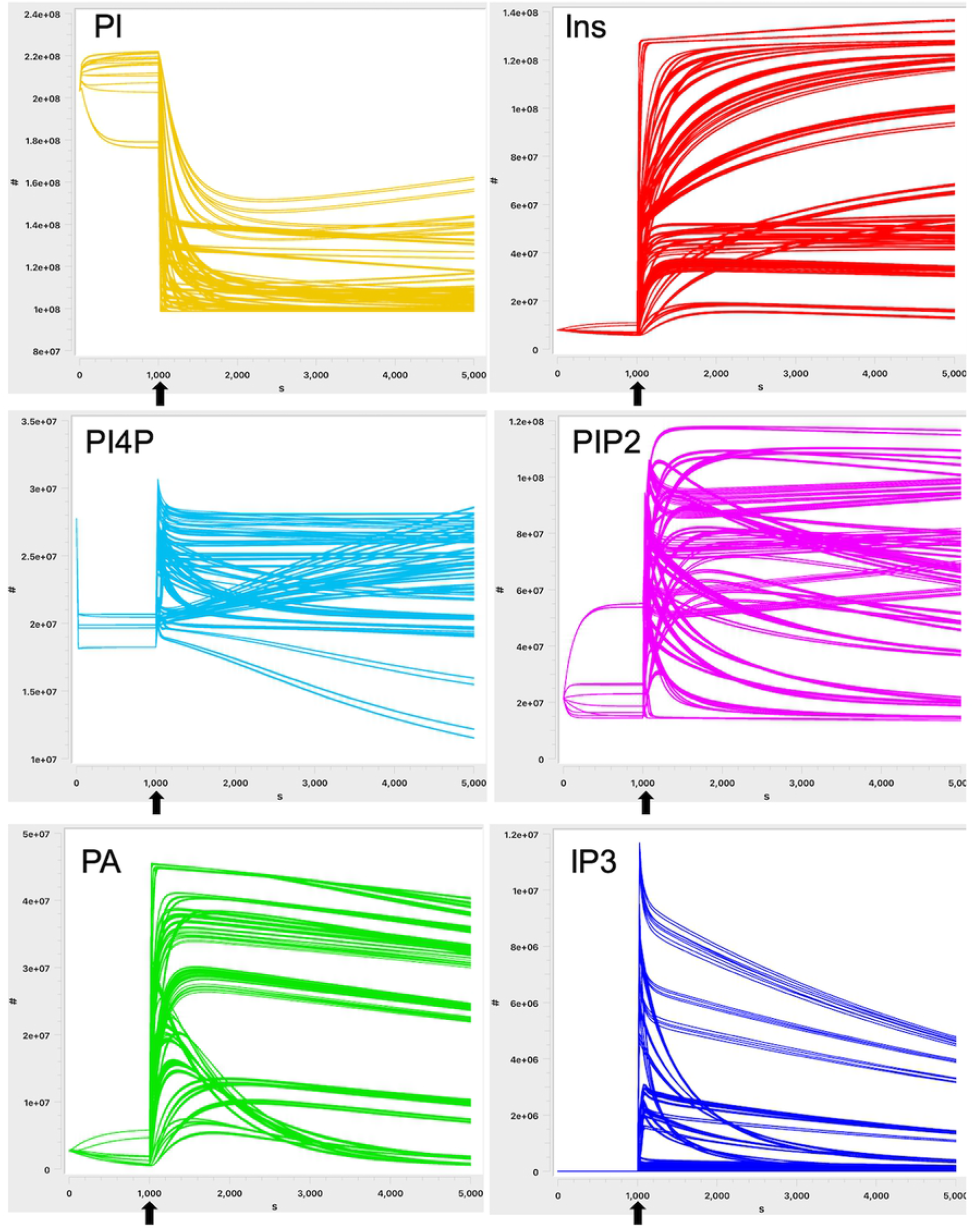
Simulation results of systematic “mix-and-match” of protein numbers between HeLa and U2OS cells proteomic data. The copy numbers of the 12 key proteins surveyed in our model (Gαq, smG, PLCβ, the combined IP3 modifying enzymes, DGK, LPP, CDIPT, OCRL1, PI4K, PIP5K, SAC1 and cPLA2) were taken from either the HeLa or U2OS proteome (12, 34), or from the calculated numbers for U2OS missing data and leading to the simulations shown in figure S4A, and systematically mixed using Parameter Scans. 75% of the 4096 simulations obtained lead to incorrect dynamic behaviours for the outputs monitored with our model. Simulations are run for 5000 seconds with activation occurring at 1000 sec (arrows), #: molecules per cell, s: seconds.

Together these results suggest that combining data from different cell types does not necessarily lead to results that are consistent in simulating PI cycle dynamics. Indeed, it is highly likely that in the majority of cases, model simulations are unlikely to agree with experimentally observed results for specific cell types. To further test this result we undertook a sensitivity analysis to determine the influence of initial protein levels on the production of IP3. This revealed both commonalities and specific patterns for each cell type (S3B Fig). Specifically, similar changes in the concentration of PLCβ, PIP5K and OCRL1 lead to distinctive IP3 outputs in the different simulated cells.

Thus, we conclude that although our model can be populated with protein concentrations sourced from different cell types, to describe functional outputs, it can lead to erroneous conclusions.

## Discussion

Conventionally, mathematical models of cell signalling pathways have been informed by data taken from a range of cell types. This is often a result of data not being available for a specific cell type to inform all kinetic rate constants and concentrations. Here we developed a biological model of the PI cycle entirely based on a single quantitative proteome and multiple sets of experimental data generated under the same conditions for a single cell type, the human platelet. This allowed us to focus on determining in situ kinetic rate parameters of the reactions governed by Gαq-coupled receptors. In addition, and in contrast to most other published cell signalling models, we have considered how the system behaviour varies from an inactive to active state. This has allowed us to reveal a number of mechanisms involved in the maintenance of the observed steady-state before activation.

Previous mathematical models have largely ignored the cyclic generation of PI, assuming instead that it was available at all times [1–3]. We postulated that signalling events lead to the depletion of PI on the plasma membrane and termination of the signalling, while leaving a pool of PI on ER and Golgi membranes. Using our model, we were able to show that while there is a constant exchange of PI between the different membranes, the rate of plasma membrane PI replenishment does not seem to be modified by activation events. This leads to the conclusion that PI distribution on the different cell membranes, and the rate at which it can be replenished at the plasma membrane, is a major way of regulating the duration of cell signalling.

While it has been suggested that sequestering of membrane lipids by binding proteins is an integral part of cell homeostasis and activation [18,18, 29,30], the involvement of PL-binding proteins in cell signalling regulation is, however, usually considered only after cell stimulation. We demonstrated that PL-binding proteins also play a stabilising role prior to signalling events, functioning as cellular sinks and limiting the availability of lipids to modifying enzymes such as PLC and PI3K; effectively inhibiting signalling processes. After signalling is triggered, PL-binding proteins also regulate the maximum amount of available PL for reactions leading to intermediates such as IP3 and DAG. Simulations of PI45P2-binding proteins correspond to the peak quantity of PI45P2 monitored experimentally and we predict that this is likely to be true for all other PL-binding proteins.

Next, our model, formulated using thrombin-stimulated platelet data, was able to recapitulate known IP3 outputs for other GPCR ligands by simply replacing the receptor numbers for each ligand with their respective known values. These results support the still-debated hypothesis that the number of GPCRs, rather than their molecular identity, regulates the intensity of this type of signalling [1,23].

We also used our model to produce a series of predictions regarding the regulation of the major enzymes involved in the central PI45P2 -PI4P -PI axis of the PI cycle. We predicted that three out of the four enzymes, namely PI4K, PIP5K and OCRL1 have their activities up- and down-regulated in concert. The co-location of PI4K and PIP5K has already been shown to occur experimentally [19] and we suggest adding OCRL1 to this complex. In contrast, to explain the differential abundancy of PI and PI4P/PI45P2 as well as the distinct response of the phosphatase SAC1 to the activation signal, we predict this enzyme to be isolated from its counterparts. SAC1 has been shown to be restricted to discrete regions of contact between the plasma and ER membranes [31]. Interestingly, CDPIT, otherwise known as PI-synthase, is also shows a lack of up-regulation after signalling [32], and is known to be located at cell membrane contact points between the ER and plasma membranes. It is likely that the PI4K-PIP5K-OCRL1 complex is recruited by either the receptors or some of their downstream effectors. There is no physical connection between the location of the complex and the membrane contact regions containing the different enzymes regulating the regeneration of PI and its location on the different membranes.

Due to the lack of detailed protein concentration and reaction rate constant values for individual cell types, the use of data obtained from multiple cell types is common in cell signalling models. Kinetic rates are often sourced from experiments performed with purified enzymes or adapted from previous modelling attempts. While mathematical models informed using ‘mosaic data’ have been crucial in understanding mechanisms that underlie processes within cells, combining data from different mathematical models of the same pathway are often difficult. This is because model formulations often differ, meaning parameter dimensions do as well. This is further compounded by the fact that data is not always available to inform all model parameters, meaning uninformed cell specific parameters are often determined using similar processes in other cell types or are simply estimated. We hypothesised that fully informing a mathematical model of a pathway using cell type-specific data would lead to more accurate predictions of the pathway dynamics. We demonstrated that our PI model of a human platelet demonstrated similar behaviour when informed by mouse platelet data, where the kinetics were assumed the same, but protein concentrations varied. In extending the model to that applicable to a nucleated HeLa mammalian cell, similar observations were made. However, maintaining the same kinetic rate constant values but using protein concentrations obtained from two different cell types led to widely varying dynamical predictions for PL and IP3.

The long-term goal of biological modelling is to recapitulate processes that govern cell behaviour. Our experience in modelling the PI cycle was that there was too much variability in kinetic reaction rate constants and protein concentration published from prior partial models to build a comprehensive model of the pathway. In order for parameterisation of mathematical models for specific cell-types to be more consistent, the collection of comprehensive experimental data informing the concentration of cellular components and their dynamic behaviour needs to occur. Currently, such values are generally informed by traditional methodologies such as western blotting or lipid chromatography, both of which are limited in the number of molecules they can simultaneously characterise. They also suffer from a lack of uniformity and reproducibility. Omics technologies are now available to process high numbers of molecules using highly standardised protocols, which should allow for such issues to be overcome [33]. Extensive quantitative proteomic, metabolomic and lipidomic datasets for commonly available cell lines already provide a valuable resource for modellers. Given the number of signalling pathways in eukaryotic cells that are being considered for mathematical modelling, the rapid expansion of cell-specific omics generated data sets, are likely to become a central part of ensuring mathematical models of signalling pathways are quantitatively well informed.

## Methods

Further details of the biological rationale and of the description of the different iterations of the model are available in the Supplementary Information document.

### Estimation of compartment sizes

Our core model consists of 3 compartments: the plasma membrane, the cytosol and the organelles. The volume occupied by the platelet plasma membrane, including the open canicular system (OCS) inside the cytoplasm, was calculated to be around 1 fl. Based on published observations, the size of the cytoplasm was reduced as the observed volume of platelets is filled by organelles such as vesicles, ER remnant, mitochondria and the OCS estimated to occupy up to 40% of the internal volume. Most of the remaining volume contains cytoskeletal proteins and glycogen granules and is calculated to occupy around 1 to 2 fl [35,38,39]. We estimated the mouse platelet plasma membrane and cytosol volumes to be around 0.25 fl each while the nucleated cells average plasma membrane and reaction cytosol volumes at around 17 fl each based on a 2000 fl overall cell volume.

### Initial parameters

Our model takes into account the concentration of protein and lipids relevant for each reaction. Data for protein copy numbers were obtained from a human platelet proteome [5] (S2 and S3 Tables) while initial and post-activation time-resolved data for the Phospholipids (PL) and Inositol Phosphates (IP) were collated from several publications [6–10,14,16,24,40–45] (S1 Table, Fig 3, S1 Fig). For collated pools of proteins, the UniProt database was first mined using the PL names, followed by a search of the quantitative proteome using the UniProt codes (S2 and S3 Tables). Although their binding affinities for the different PL are likely to be variable, we parametised the different on/off rates on the assumption of average kinetics.

### Estimating kinetic rate constants

We used the free software COPASI v4 [15] to build our model starting from a reduced basic network of reactions. All molecules were considered as “well-mixed” inside each compartment. Mass Action kinetics were assumed in modelling the respective reactions. The governing equations were solved using the deterministic method (LSODA) solver. Reaction enzymes were not written as modifiers as their concentrations were important for our model. Activation by ligands were written as events after the start of the simulation. All enzymatic activations were terminated by either the production of an inactive protein (inactivation) or by the return to the initial basal activity state (reset). For each reaction the default rate was first selected then a series of parameter scans were performed, starting at +/− 4 orders of magnitude until the values were producing time-resolved curves for each and every PL and IP output in the model matching the experimental data using a fit-by-eye. Steady-state analysis and Time Series Sensitivity Analysis were performed on all reactions that occur before the activation event, and the protein and PL/IP concentrations adapted accordingly. Parameters sets that led to a deviation of more than 20% from the experimental data were rejected and the full analysis restarted. Schematic diagrams of the reactions and the parameters are shown in Table S7. The computed initial molecule numbers are listed in Table S6. The model was deposited in the BioModel database [46]

## Acknowledgments

We thank Dr J. Rudge and Prof P. Dash for useful discussions and advice during the development of the model. This work was funded by the British Heart Foundation (PG/16/20/32074).

## Supplementary Figure legends

**S1 Fig.** Core model development

A: Collated graphs of experimental data used to inform the model, adapted from (5–10, 22). See Table S1 for details and Fig 2 for standard deviations. Activation occurs at t=0 sec. PIP3 and PI34P2 results are shown but were not used in our model. B: Summary diagram of Model Iteration 1 reactions. Only key lipids were kept namely PI and PI45P2, Inositol (Ins) and Phosphatidic Acid (PA). The reactions in this diagram describe a PI cycle in a non-activated cell. C: Summary diagram of the Gαq-protein coupled receptor activation cascade. See Table S5 for details and parameters. D: Summary diagram of Model 1 Iteration reactions in an activated cell. Reactions rates k_2_ and k_3_ are now replaced by k_2_’ and k_3_’ representing the activated rates of the respective enzymes. Coincidence detection leading to the change of rate k_3_ in k_3_’ is shown in green. The activation of cPLA2 leads to the removal of some of the PA from the plasma membrane to produce arachidonic acid (k_5_). k_1_ and k_4_ remain unchanged. E: Summary diagram of the addition of lipid binding proteins to regulate the availability of the phospholipids. Although only the activated state is shown in this diagram, the binding and release of the PL was assumed to be constant under both inactivated and activated states of the cell. F: Summary diagram showing the final iteration of the PI cycle model including the addition of PI4P, DAG and IP3, their respective modifying enzymes and binding proteins. Inactivation reactions not shown. See Table S5 for details and parameters.

**S2 Fig. Description of the first iterations of the core model.** A: Graphical summary of the reactions in the early iterations for either inactivated or activated states. Activated GPCRs (RGa) trigger the activation of the PIP5-kinase (PIP5Ka) together with PA. The levels of PA in the cells are also regulated by activated cPLA2 (PLA2a) which produces arachidonic acid (AA), a precursor of Prostaglandin H2 (itself a precursor of TxA2 produced by the platelet as a secondary signaling molecule) and several eicosanoids. B: Table summarising the different steps in the early core model construction and the output for PI45P2 and PA. C-D: Graphs of the results shown in the table for the simulations 1A-B. E: Graphs of the results shown in the table for each simulation 1C., the results were virtually identical whether the activation of PIP5K was simulated directly via the activated GPCR or indirectly by the activated PLC (PLCa). Activation time points are indicated by arrows.

**S3 Fig. Additional analyses of the Core Model in other cell types.** A: Comparison of the core model outputs in human and mouse platelets. The volume of the mouse platelet, PL and IP initial concentrations have been modified as described in the material and methods. The numbers of GPCR receptors for each platelet type are listed in Table S4. The overall results for the PL, Ins and IP3 are virtually identical except for the overall levels which are related to the initial amounts in the two cell types. The simulations have been extended to 10000 seconds to capture any late trend, with the activation occurring at 1000 sec (arrows). B: Comparison of Sensitivity Analyses for IP3 in nucleated cells simulations. Time series Sensitivity Analyses of the impact of some key protein initial concentrations on IP3 outputs were performed and compared when our model was populated with either human platelet, HeLa or U2OS proteomic data. We used protein numbers estimated via Parameter Scans for missing protein values in the U2OS proteomic dataset namely Gαq, cPLA2, PI4K, OCRL1 and IP3 modifying enzymes (IP3E). PIS = CDIPT. IP3 results for each protein initial concentration are shown in the table.

**S1 Table:** Summary table of published Phospholipids and Inositol Phosphates experimental data.

**S2 Table:** Quantification of Lipid binding Proteins in human platelets.

**S3 Table:** Relevant protein number and UniProt codes.

**S4 Table:** A: Protein numbers in HeLa, U2OS, mouse platelets, compared to human platelets. All data from respective proteome datasets unless stated otherwise. B: reaction volumes and Gαq-coupled receptor numbers for each cell type simulations.

**S5 Table:** Schematic diagrams of the reactions and parameters.

**S6 Table:** Initial particle number and concentrations for human.

